# Improved structural variant discovery in hard-to-call regions using sample-specific string detection from accurate long reads

**DOI:** 10.1101/2022.02.12.480198

**Authors:** Luca Denti, Parsoa Khorsand, Paola Bonizzoni, Fereydoun Hormozdiari, Rayan Chikhi

## Abstract

Structural variants (SVs) account for a large amount of sequence variability across genomes and play an important role in human genomics and precision medicine. Despite intense efforts over the years, the discovery of SVs in individuals remains challenging due to the diploid and highly repetitive structure of the human genome, and by the presence of SVs that vastly exceed sequencing read lengths. However, the recent introduction of low-error long-read sequencing technologies such as PacBio HiFi may finally enable to overcome these barriers. Here we present `SVDSS`, a novel hybrid method for discovery of SVs from long-read sequencing technologies (e.g., PacBio HiFi) that combines and effectively leverages mapping-free, mapping-based and assembly-based methodologies for overall superior SV discovery performance. Our experiments on several human samples show that `SVDSS` outperforms state-of-the-art mapping-based methods for discovery of insertion and deletion SVs in PacBio HiFi reads and achieves significant improvements in calling SVs in repetitive regions of the genome.

`SVDSS` is open source and publicly available at: https://github.com/Parsoa/SVDSS

## 1 Introduction

Structural variants (SVs) are defined as medium to large-size genomic rearrangements [1, 2]. SVs can range from tens of base-pairs to over megabases of sequence. The different types of SVs include balanced SVs, such as inversions and translocations, and unbalanced SVs, such as insertions and deletions [3]. The study and characterization of SVs has been driven by constant improvements in the available technologies to assay variants. Although structural variants are not the most ubiquitous type of genetic variants, the total volume of base-pairs impacted by SVs is far more than any other type of genetic variant, including Single Nucleotide Variants (SNVs) [4, 5]. Furthermore, recent studies of structural variants using orthogonal technologies has shown that SVs are the least well-characterized type of genetic variants with many basic questions still not completely resolved, such as the average number of SVs per sample or sequence biases contributing to their formation [6, 7, 8, 9]. In addition, the homology-driven mechanisms behind SV formation (e.g., non-allelic homologous recombination) has contributed to the complexity of their systematic study [10]. It is believed that a large fraction of polymorphic SVs are still not fully characterized [11, 12].

As our current understanding of SVs evolves, it is becoming clear that SVs are a major contributing factor to human diseases [13, 14, 15], population genomics [5, 16] and evolution [17]. The comparative study of SVs in multiple closely-related species (e.g., great apes) has shown significant contribution of SVs to evolution (e.g., through gene duplication or deletion [18, 19]). Furthermore, study of rare and *de novo* SVs in disease such as autism and epilepsy has proven the significant contribution of these variants in such diseases [20, 21, 22, 23, 15]. It is also known that somatic SVs are one of the major causative variants in different types of cancer [24, 25, 26, 27].

With the advent of short-read and long-read high-throughput sequencing technologies in the past decade, significant progress has been made in our understanding of the abundance, complexity, and importance of SVs [28, 29, 30, 31]. There are many methods developed for prediction of SVs using whole-genome sequencing (WGS) data produced from different sequencing technologies [32, 33, 34, 35, 36, 37, 38, 39, 40, 41, 35, 42]. Majority of these methods try to predict variants by detecting certain SV signatures (i.e., read-depth, read-pair, or split-read) in mappings of the reads to the reference genome [43, 44, 6] and are hence known as “mapping-based” methods. Mapping-based methods have contributed significantly to our understanding of the abundance of SVs in the general population and their role in diseases [11, 45, 46]. Mapping-free methods are a more recent group of approaches that try to predict SVs without mapping the reads to the reference genome and instead by comparing sequence data between different genomes [47, 48]. Finally, assembly-based approaches first assemble the sequenced reads into longer contigs and use the assembled contigs to predict variants [49, 50, 51]. Assembly-based methods have recently been shown to provide superior performance to mapping-based tools [52].

There are several limiting factors for predicting SVs using each of these frameworks. First, since majority of SV prediction tools use mappings of the reads to the reference genome for making SV calls, predicting SVs in highly repeated regions of the genome (e.g., segmental duplications) where mappings can be inaccurate would be prone to false discovery. Furthermore, predicting complex SVs such as inversion-duplications - that account for a significant fraction of SVs - using purely mapping-based approaches can result in increased false discovery rates compared to basic SV types [38, 53, 36, 8, 54]. Finally, the reference genome gaps and misassemblies further complicate the prediction of SVs in these regions and result in decreased accuracy and increased variability across tools [8]. The mapping-free approaches on the other hand suffer from not being able to provide the loci of the event. Furthermore, the direct sequence (or the fixed-length *k* -mer) comparison performed in these type of tools can result in collapse of repeats and lower sensitivity/accuracy. Finally, assembly-based approaches are very computationally resource intensive and often require integration of data from multiple different technologies (i.e., long-reads, short-reads, and Hi-C) [55, 52], higher sequencing depths (35x was reported in [52]), and extensive polishing and post-processing to yield a high-quality *de novo* assembly suitable for variant prediction, thus making them impractical for large-scale SV discovery across many samples. Furthermore, these approaches usually represent a single hapoltype and might miss variants in the diploid genomes [56].

Here, we propose a novel method called `SVDSS` that combines advantages of all three mapping-based, mapping-free, and assembly-based approaches for predicting SVs. Our method utilizes mapping-free sample-specific signatures [57] along with mapping information to cluster reads potentially including SVs and then performs local assembly and alignment of the clusters for SV prediction. With the combination of different analysis methods, our algorithm is able to improve SV calling performance particularly in repetitive areas of the genome compared to other contemporary approaches.

## 2 Methods

We present `SVDSS` (Structural Variant Discovery with Sample-specific Strings), a novel method for the discovery of structural variants from accurate long reads (e.g., PacBio HiFi). `SVDSS` takes as input a reference genome and a mapped BAM file and produces SV calls in VCF format along with assembled contigs for SV sites in SAM format. We use the concept of sample-specific strings (SFS) which we previously introduced as the shortest substrings unique to one string set with regards to another string set [57]. We employ SFS here to pinpoint differences between reads and a reference genome [57]. Our method utilizes SFS extracted using an optimal algorithm [57] for coarse-grained identification of potential SV sites. It then performs Partial Order Alignment (POA) [58] of clusters of SFS from such sites to produce contigs that are then locally aligned to the reference genome to detect SVs. The main advantage of using SFS is that they are not limited to fixed length seeds (unlike *k*-mers) and the algorithm can dynamically find the shortest string for covering the breakpoints of each variant, thus, making SFS ideal for anchoring potential SV breakpoints.

`SVDSS` has three main steps as depicted in Figure 1, sketched here and explained in more details in the following sections:

**Figure 1:**
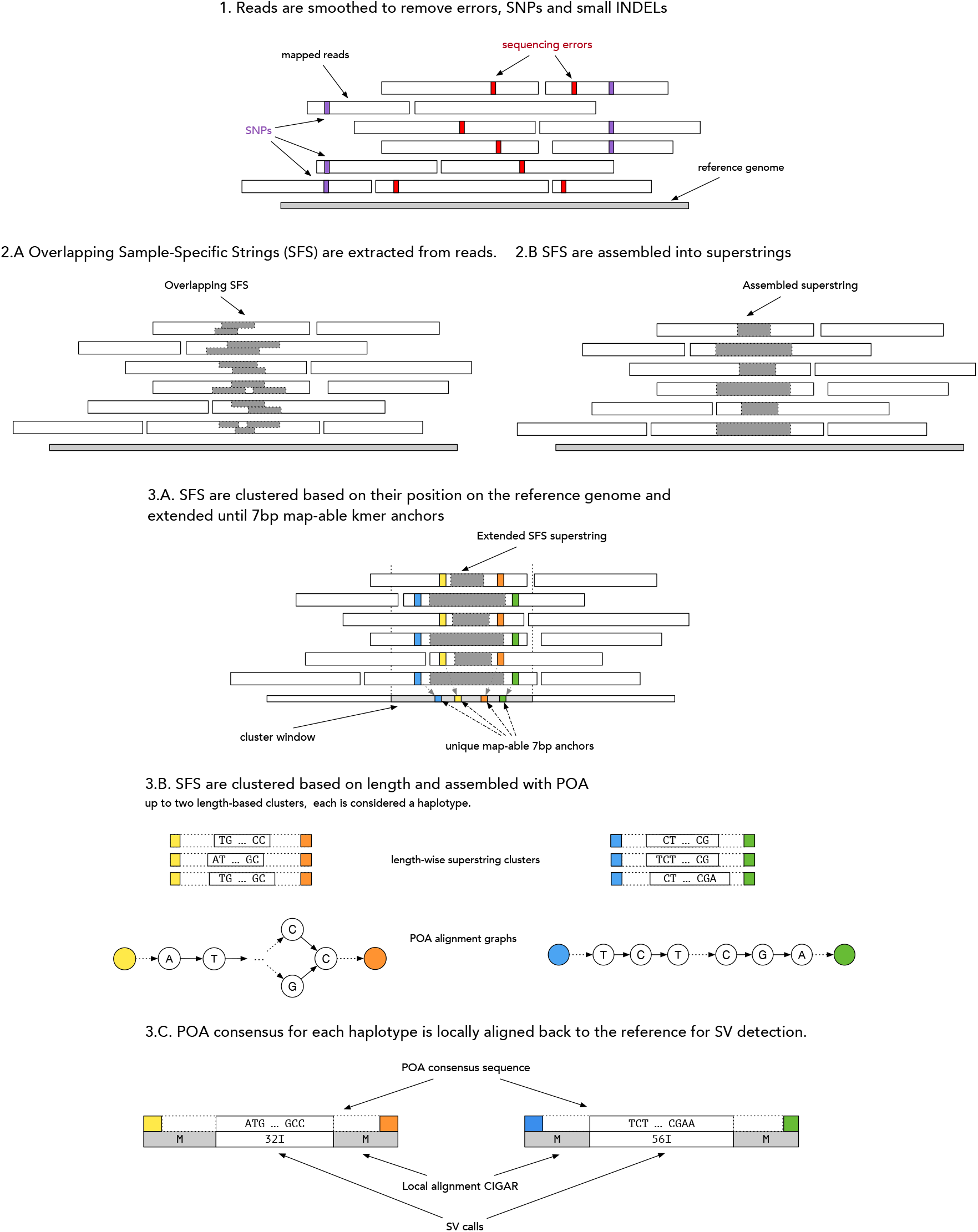
Overview of the SVDSS SV prediction pipeline. 1) Reads are smoothed to remove SNPs and sequenceing errors. 2) SFS are extracted from reads (A) and assembled into superstrings (B). 3. Superstrings (grey) are clustered based on their placements on the reference genome and extended until uniquely mapable 7bp anchors on each side (colored) (A). Each cluster is further clustered into up to two subclusters based on length of the superstring. Each subcluster signifies a potential haplotype. The subclusters are assembled with POA to generate a consensus sequence (B). The POA consensus for each cluster is locally aligned to the reference genome and SVs are called from the mapping information.

1. **Read smoothing:** reads are modified to remove sequencing errors, SNPs and small indels (*<* 20bp) that may interfere with SV calling (Figure 1 step 1, and Supplementary Figure S1). Smoothing significantly reduces the number of extracted SFS while increasing their specificity for the purpose of SV calling.
2. **SFS superstring construction:** SFS are computed from the smoothed reads using the optimal Ping-Pong algorithm [57] (Figure 1, 2A) and then assembled into superstrings to reduce redundancy (2B).
3. **SV prediction using SFS superstrings:** SFS superstrings are clustered based on position and extended to include unique anchoring sequences from the reference genome (Figure 1, 3A), further subclustered by length then assembled based on POA approach to generate haplotype candidates (3B). Finally SVs are called by aligning the resulting POA consensus(es) (3C).

**Figure 2:**
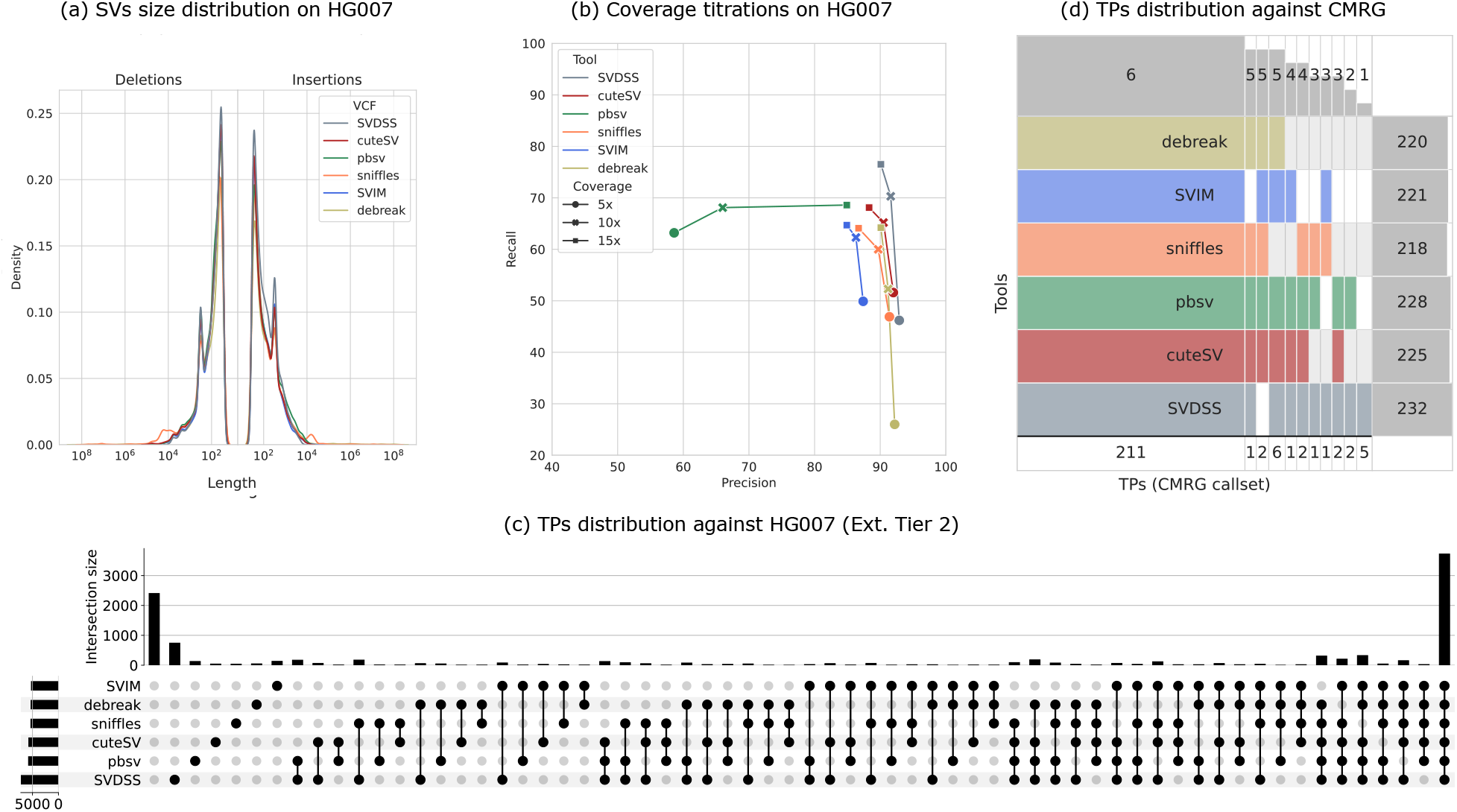
(a) Distribution of SVs lengths reported by different tools on HG007 (Full Genome). (b) Lineplot presenting results of the coverage titration for 5x, 10x, and 15x. (b) Analysis of shared calls (True Positives) between different tools on HG007 (extended tier 2). (d) SuperVenn diagram showing shared calls (True Positives) between different tools on the 273 medically-relevant genes considered in the CMRG callset.

**Figure 3:**
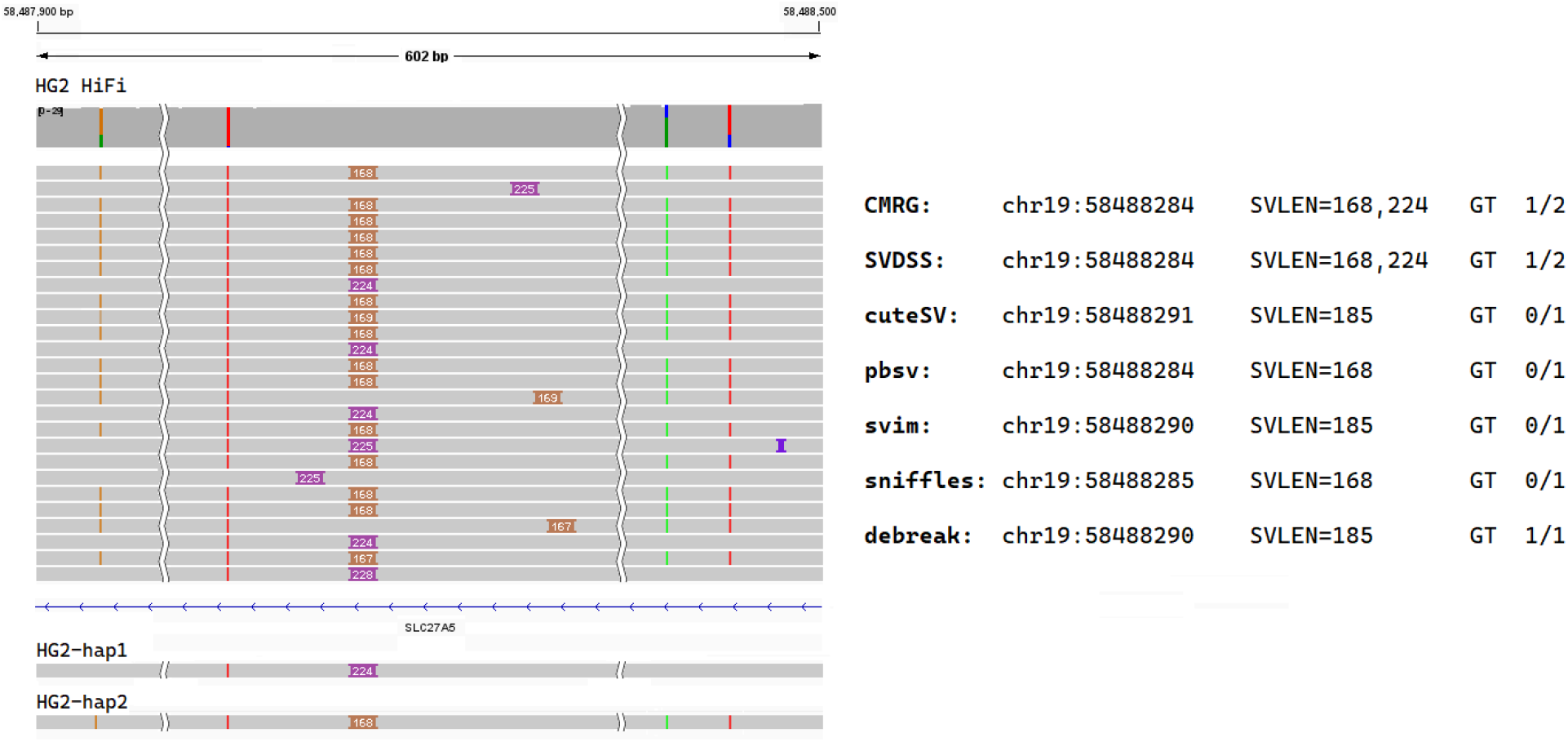
Example of an SV at a medically-relevant gene that has been correctly called exclusively by SVDSS. On the right panel, IGV sketch of the 602bp region around the SV (the full region is reported in Supplementary Figure S10). The sketch reports the HiFi reads alignment along with the haplotype alignment performed using `minimap2` (as part of the `dipcall` pipeline). On the left panel, details on the SVs reported by the CMRG callset, `SVDSS`, and the other alignment-based callers considered in our evaluation.

### 2.1 Sample-specific string computation and assembly

Sample-specific Substring-free Strings (SFS) are defined as sequences that are specific to a “target” set of strings (a genome or sequencing sample) with respect to another “reference” set of strings (another genome or sequencing sample) [57]. The “substring-free” part means that they do not occur as substrings of each other. Note that, in the context of SV discovery, the “reference” will always be an assembled reference genome, e.g., GRCh38, and the “target” here is a set of reads. SFS can be optimally computed using the Ping-Pong algorithm, presented in [57]. Ping-Pong builds the FMD index [59] of the reference genome and queries the reads of the target sample against this index to report substrings that are not present in the index. The FMD index is a bidirectional text index with constant-time forward and backward search operations, thus allowing for efficient computation of SFS.

When SFS are computed between a reference genome and a target sample, they capture nearly all variations expressed in the sample with respect to the reference genome, as shown in [57]. Indeed, each sequencing read including a variant produces at least one SFS supporting the variant, hence a variant will be supported by at least one SFS per read covering it. SV breakpoints usually result in novel sequences that are captured as SFS. However, due to the “shortest” property of SFS, the entire SV sequence is not necessarily covered by a single SFS: a read may produce several overlapping SFS for long variations. To remove unnecessary redundancy in the information captured by overlapping SFS, we newly assemble all such overlapping SFS into longer strings called “superstrings”. Assembling SFS into superstrings also reduces the number of SFS by an order of magnitude, making any downstream analysis more efficient.

As SFS on each read are naturally sorted based on their start positions, the assembly stage can be implemented as a single pass over the SFS on each read, merging each SFS with the next one if they overlap. The resulting superstring can further be merged with the next SFS if they also overlap, etc. More formally, on a read *R* where *k* consecutive SFS are overlapping such that *R*[*i*_1_, *j*_1_] overlaps with *R*[*i*_2_, *j*_2_] and *R*[*i*_2_, *j*_2_] overlaps with *R*[*i*_3_, *j*_3_] and … *R*[*i*_*k−*1_, *j*_*k−*1_] overlaps with *R*[*i*_*k*_, *j*_*k*_], we merge the strings into the single superstring *R*[*i*_1_, *j*_*k*_].

The SFS assembly procedure effectively merges all the SFS belonging to the same variant into a single long superstring. This results in superstrings from the same variant to have similar length, sequence and position with respect to the reference genome which allows them to be easily clustered for SV prediction.

### 2.2 Read Smoothing

The SFS extraction step (Ping-Pong algorithm) requires reads with low error-rates for optimal performance as sequencing errors can result in millions of undesirable SFS. While most such SFS can be filtered later on, they can negatively affect the accuracy and will increase runtime by adding excess processing. Furthermore, the presence of millions of SNPs and small indels in a sample also results in tens of millions of additional SFS being extracted that are not directly useful for genotyping SVs. To solve both of the above problems, we introduce a preprocessing step called *“read smoothing”* that aims to eliminate both sequencing errors and short variants from input reads. The smoothing algorithm starts from read alignments (a BAM file) and uses information from the CIGAR strings of each alignment to remove any short mismatch between a read and the reference genome.

In more details, for segments reported as a match between a read and the reference genome (CIGAR operation `‘M’`), the algorithm replaces the read sequence with the corresponding sequence from the reference genome, automatically removing any single-base mismatches (i.e., sequencing errors or potential SNPs) in the process. For short deletions (CIGAR operation `‘D’`), the algorithm removes the deletion from the read by copying back the deleted bases from the reference sequence. Short insertions (CIGAR operation `‘I’`) are similarly smoothed by removing the inserted bases from the read. Using the default parameters, deletions and insertions are smoothed if they are shorter than 20bp. Note that smoothing insertions or deletions, i.e., removing them from the alignment, results in the extension of the `‘M’` sections of the CIGAR string. Finally, soft-clipped regions (CIGAR operation `‘S’`) are retained as they include potentially long inserted or deleted sequences: any SNP or sequencing error inside clipped regions cannot be corrected as a result. As a result of the smoothing algorithm, a smoothed read’s CIGAR strings will have significantly fewer edit operations than that the original read and it will consist of one or more very long `‘M’` segments with large INDELs in between, potentially surrounded with soft-clipped regions. Supplementary Figure S1 illustrates the smoothing procedure on an example read. We note that the Ping-Pong algorithm will not produce any SFS that is entirely contained in a M section of a smoothed read as the corresponding sequence has been replaced base-by-base with reference genome sequence. Therefore, the number of SFS extracted from smoothed reads is significantly smaller than the number of SFS extracted from original reads.

Smoothing relies on correctness of read alignments. If an alignment is thought to be inaccurate, the smoothing algorithm does not modify it. To this aim, during its execution, the algorithm keeps track of the average number of mismatches between the `‘M’` segments of alignments and the corresponding reference sequence: any read that has more than 3 times the average mismatch rate is ignored, i.e., is not modified.

On a more technical note, we point out that the above modifications do not change the overall mapping of the read as the mapping positions (begin and end) remain the same. As a result, the algorithm will not change the order of the reads in a sorted BAM file. This allows us to quickly reconstruct a sorted BAM file without the need to sort it again. However, because the size of the reads may have changed, the index of the original BAM files is no longer valid for the smoothed BAM and it has to be indexed again with `samtools index`.

In our experiments, smoothing effectively reduces the number of extracted SFS by over 90%, while having effectively no impact on the SV calling pipeline’s recall. Out of the 6.2M reads for the CHM13 samples, around 5M are smoothed and the rest are deemed to have unreliable mappings and are discarded. The 1.2M non-smoothed reads from CHM13 are responsible for more than 82% of all SFS extracted from that sample after smoothing. However, the SFS extracted from non-smoothed reads do not contribute to increasing the method’s recall at all. Indeed excluding the SFS extracted from non-smoothed reads increases the method’s precision while leaving the recall unaffected. This justifies the exclusion of non-smoothed reads from the `SVDSS` pipeline. Further analysis shows that nearly all non-smoothed reads map to centromere regions of the CHM13. Supplementary Figure S2 shows the distribution of mapping positions of reads from chr1 on both CHM13 and GRCh38. The large gap around the centromere when mapping to GRCh38 explains the poor performance of non-smoothed reads when predicting SVs against the reference genome.

In summary, read smoothing is a critical preprocessing step of the `SVDSS` pipeline. It reduces the number of retrieved SFS and increases the specificity of the extracted SFS which results in higher precision in predicting SVs without deteriorating recall. The procedure is also computationally very lightweight, as it essentially rewrites the BAM file in a single pass with minor modifications. As a result, smoothing is an effective method for increasing the specificity of SFS for SV calling and improving the computational efficiency of the pipeline.

### 2.3 SV Discovery

The main SV calling algorithm consists of three main steps (see Figure 1 steps 3A, 3B, 3C):

1. Superstrings constructed from the SFS strings are “placed” on the reference genome by extracting their alignments from read alignments. The superstrings are then clustered based on their aligned loci. Each cluster represents one or more SVs that are close to each other and may also contain multiple alleles.
2. Each cluster is further clustered based on length to generate up to two haplotype candidates (taking into account the diploidy of the human genome). Each haplotype cluster candidate is assembled with Partial Order Alignment (POA) to yield a consensus sequence.
3. Each haplotype candidate is locally realigned back to the reference genome region corresponding to its cluster and SVs are called based on the alignment.

We will explain each step in more details in the following subsections.

#### 2.3.1 Superstring placement and clustering

Aligning superstrings back to the reference genome can be time-consuming and error-prone due to their relatively short lengths. However we note that the necessary mapping information is indeed already available in the sample’s read mappings. In practice we do not align superstrings directly to the reference genome but instead their alignment is extracted from the alignment of their originating reads. Assuming *R*[*i, j*] is a superstring that spans positions *i* … *j* on read *R*, by knowing the mapping position of *R*, we can easily place the superstring on the reference genome by analyzing the corresponding CIGAR portion (i.e., CIGAR sections covering positions *i* … *j*).

It is possible that a superstring’s mapping is entirely contained in an inserted or clipped part of the read and hence the mapping extracted above cannot be used to place the superstring on the reference genome. To avoid this, each superstring is extended on the read on both sides until we reach a perfectly mappable (i.e., that can be map to the reference genome with no errors) and locally unique (i.e., is not repeated in the considered window) *k*-mer anchor. The default value for *k* is 7 and the default window size is 100bp (on each side of the superstring). The superstrings that cannot be extended in this manner are ignored. Figure 1 3A shows this extension procedure. The *k*-mer anchoring idea was influenced by LongShot [60].

Finally, we cluster the superstrings based on their mapping locations: superstrings that have close enough mappings (by default less than 500bp apart) are placed in the same cluster. The resulting cluster’s interval is defined as the smallest interval in the genome that completely includes all of its superstrings and the includes either a single SV or several close or overlapping SVs possibly from different haplotypes.

#### 2.3.2 Cluster POA and SV detection

Each cluster so far includes one or more close SVs. However, as the human genome is diploid, the SVs might indeed be from different haplotypes. To resolve the different haplotypes, we further split each cluster into subclusters of superstrings of similar size and sequence. This is based on the assumption that different alleles at each site have different length and sequence. The similarity of sequences is calculated using `rapidfuzz` ^1^. The two largest resulting subclusters (in terms of number of superstrings) are selected as haplotype candidates (considering the human genome is diploid). If only one subcluster is returned, it signifies a homozygous variant. `SVDSS` then computes a consensus sequence for each subcluster using Partial Order Alignment.

Assume that a cluster *c* spans the interval *G*[*s*_*c*_, *e*_*c*_] of the reference genome *G*. Most strings of the cluster only partially cover this interval (i.e., they align to positions [*s, e*] with *s*_*c*_ *≤ s < e ≤ e*_*c*_) while some others span the entire interval (i.e., they align to positions covering at least [*s*_*c*_, *e*_*c*_]). In order to perform a more accurate POA, `SVDSS` requires all the strings in a cluster to be of the same length. Therefore, `SVDSS` fills the gaps preceding or following a superstring using the reference genome. For instance, if a superstring *S* aligns to [*s, e*] with *s*_*c*_ *< s < e < e*_*c*_, then the resulting sequence will be *G*[*s*_*G*_, *s* 1] + *S* + *G*[*e* + 1, *e*_*G*_] (where + is the string concatenation operator). The main goal of this extension is to summarize the information contained in a cluster and to minimize the difference between the superstrings coming from different reads. The extended superstrings in each subcluster are then aligned to each other using `abPOA` [61] to generate a consensus (Figure 1 3B).

Finally, each POA consensus sequence is realigned locally to the reference genome window corresponding to its cluster using `parasail` [62] and the alignment’s CIGAR information is analyzed to call and detect SVs (Figure 1 3C). A weight is assigned to each SV prediction based on the number of superstrings that support it. A higher support indicates a more confident call. By default, we filter out SV calls having less than 4 supporting superstrings. The confidence threshold can also be set at runtime using the `--min-cluster-weight`option.

#### 2.3.3 SV Chain Filtering

In repetitive regions of the genome such as STRs, the nature of repeats may result in reads originating from the same locus to map to slightly different coordinates. This will result in multiple clusters (relatively close to each other) and multiple SV calls for the same variant but at slightly different positions. To reduce the number of false positives and eliminate such redundant calls, we perform a “chain-filtering” post-processing step. This step sorts all predicted SVs based on coordinates and filters out consecutive SVs of the same type with similar sizes, keeping only the one with the highest weight.

### 2.4 Implementation details

As a result of its many steps, `SVDSS` is more compute-intensive than other SV discovery methods, yet remains practical to run. In this section we comment on the performance of each of the steps. Our implementation has been optimized to utilize multiple-core processors. The FMD-index creation and querying are handled internally by the FMD implementation from [59]. FMD-index creation for the GRC38 reference genome takes around 30 minutes on 16 cores. The index can be reused for any number of samples so its creation is a one-time expense.

Read smoothing is an IO-intensive step and benefits significantly from enabling the multithreaded BAM decoding functionality built into `htslib` [63] by setting the `bgzf_mt` flag when opening a BAM file. To further improve decompression performance, we require that `htslib` is built with `libdeflate` ^2^ in place of the default BAM decoder. For HiFi data at 30x coverage the smoothing algorithm takes about 15 minutes to run on 16 cores.

SFS extraction is the most computationally intensive step and takes about 45 minutes on 16 threads for the CHM13 HiFi data. Finally the SV calling steps is fast and take less than 8 minutes. Overall the runtime of the `SVDSS` pipeline is less than 2 hours for a high-coverage HiFi sample on 16 cores.

## 3 Results

In the following sections, using experimental analysis on multiple WGS samples, we demonstrate that `SVDSS` accurately predicts SVs and outperforms state-of-the-art approaches. We further show that the main contribution of our proposed approach is our ability to more accurately predict SVs falling in repeated regions of the genome compared to other methods.

### Benchmark and evaluation callsets

One complexity in comparing different tools for calling SVs is the imperfectness of available callsets. Missing variants and potentially false predictions affect almost all published callsets, and even the most high-quality callsets have been reported to have a *∼* 5% false discovery rate and a much higher false negative rate [6]. Furthermore, many callsets are constructed using state-of-the-art but imperfect SV prediction tools and are thus biased towards these methods [64]. For these reasons, we have opted out of using pre-existing callsets such as Genome In A Bottle (GIAB) [65] in our experimental benchmarking, and instead constructed our ground truth SV callsets from scratch using high-quality *de novo* assemblies (CHM13, HG002, and HG007, described in the next paragraph). We applied the assembly-to-assembly SV calling tool `dipcall` [64] to each assembly versus the GRCh38 reference genome.

### SVDSS **enables accurate detection of insertions and deletions using accurate long reads**

We experimentally validated the accuracy of the `SVDSS` pipeline in calling SVs from three whole-genome sequenced samples sequenced using PacBio HiFi technology: the homozygous CHM13 sample from the telomore-to-telomere (T2T) project [55] and the HG002 and HG007 samples from the GIAB project [65] corrected using DeepConsensus [66]. These samples were chosen because of the availability of high-quality and effectively complete assemblies for them. Furthermore, the DeepConsensus corrected HG002 and HG007 samples show higher accuracy than standard HiFi samples corrected using only `pbccs` [66]. The use of both homozygous (CHM13) and heterozygous (HG002 and HG007) samples allows for more comprehensive analysis and comparison of SV calling methods.

We mapped each sample against the reference genome using `pbmm2` [67] and then we called SVs on each sample using the `SVDSS` pipeline. We compared our approach to five state-of-the-art mapping-based SV callers: `pbsv` [68], `cuteSV` [69], `sniffles` [70], `SVIM` [71], and a recent preprint on a POA-based method, `debreak` [72]. We ran each caller setting the minimum SV support to 4 when analyzing the 30x CHM13 sample and to 2 when analyzing the 15x HG002 and HG007 samples. We then examined their insertions and deletions calls. We validated the calls of each tool against the set of SVs constructed with `dipcall` using `Truvari` [73], a SV evaluation framework which reports precision, recall, and F1 score for each method. From this comparison, we further exclude calls made in regions of the reference genome not covered by both haplotypes, as any such call would be classified as false positive regardless of correctness.

Our method consistently outperforms other methods in all three measures of accuracy when considering the full genome (Table 1 - *Full Genome* row). On HG002 and HG007 samples, `SVDSS` outperforms the other callers’ recall by 5-10% while achieving the highest (or the second highest) precision on the full genome. `SVDSS` has been able to report 2,342 (+10% w.r.t. second-best approach) more correct calls on HG002 and 1,631 (+8%) more calls on HG007 without introducing many false calls. `SVDSS` also achieves the highest recall on CHM13 and reports 782 (+2%) more true positive calls than other methods while maintaining a very high precision. While `SVDSS` has the highest F1 score on CHM13, we note that the whole-genome improvements achieved by `SVDSS` over other approaches is less significant for this sample compared to the other two samples (improvement of 2-5% in recall and 1% in F1 while achieving similar precision to other tools). This is likely due to the homozygous nature of CHM13 making SV calling relatively easier for all approaches.

**Table 1:** Comparison of performance of SVDSS and other methods on calling SVs. Accuracy of each tool is reported in terms of Precision (P), Recall (R), and F-measure (F1). Results are further broken down by considered regions of the genome. Tier 1 accounts for nearly 86% of the genome and includes mostly easy-to-genotype regions. Extended Tier 2 accounts for the remaining 14% of the genome and includes repetitive regions that are more difficult to genotype. See Supplementary Figure S3 for more detail on tiers.

| Region | Tool | HG002 |  |  | HG007 |  |  | CHM13 |  |  |
| --- | --- | --- | --- | --- | --- | --- | --- | --- | --- | --- |
|  |  | P | R | F1 | P | R | F1 | P | R | F1 |
| Full Genome | SVDSS | 88.4 | **78.2** | **83.0** | **90.1** | **76.5** | **82.7** | 87.3 | **84.6** | **86.0** |
|  | cuteSV | 86.0 | 68.6 | 76.3 | 88.3 | 68.1 | 76.9 | 87.1 | 79.7 | 83.2 |
|  | pbsv | 86.9 | 68.8 | 76.8 | 84.9 | 68.6 | 75.9 | 84.6 | 82.7 | 83.6 |
|  | sniffles | 82.0 | 67.3 | 73.9 | 86.7 | 64.1 | 73.7 | 86.4 | 81.4 | 83.8 |
|  | SVIM | 83.5 | 65.1 | 73.2 | 84.9 | 64.7 | 73.4 | **90.1** | 79.9 | 84.7 |
|  | debreak | **88.6** | 67.5 | 76.6 | **90.1** | 64.2 | 75.0 | 83.7 | 79.6 | 81.6 |
| Tier 1 | SVDSS | 95.2 | **85.5** | **90.1** | 95.2 | **82.7** | **88.5** | 95.3 | 93.4 | **94.5** |
|  | cuteSV | 90.9 | 82.9 | 86.7 | 93.0 | 79.9 | 86.0 | 94.8 | 93.1 | 93.9 |
|  | pbsv | 95.7 | 83.1 | 89.0 | 89.7 | 80.5 | 84.9 | 94.0 | 93.7 | 93.9 |
|  | sniffles | 87.7 | 81.1 | 84.3 | 92.3 | 75.9 | 83.3 | 87.2 | 93.6 | 90.3 |
|  | SVIM | 90.1 | 81.1 | 85.4 | 91.5 | 77.9 | 84.2 | **96.6** | 92.5 | **94.5** |
|  | debreak | **96.8** | 82.5 | 89.1 | **96.2** | 76.4 | 85.2 | 93.7 | 93.0 | 93.3 |
| Extended Tier 2 | SVDSS | **82.7** | **72.3** | **77.2** | **84.6** | **70.2** | **76.7** | 80.3 | **77.4** | **78.8** |
|  | cuteSV | 80.9 | 57.0 | 66.9 | 82.3 | 56.0 | 66.6 | 79.9 | 68.1 | 73.6 |
|  | pbsv | 78.4 | 57.2 | 66.1 | 78.8 | 56.4 | 65.7 | 76.0 | 73.3 | 74.6 |
|  | sniffles | 77.8 | 56.1 | 65.2 | 80.3 | 52.1 | 63.2 | 72.7 | 73.2 | 72.9 |
|  | SVIM | 76.4 | 52.0 | 61.9 | 76.2 | 51.2 | 61.2 | **83.4** | 69.3 | 75.7 |
|  | debreak | 80.4 | 55.3 | 65.5 | 82.3 | 51.9 | 63.7 | 74.4 | 68.1 | 71.1 |

Figure 2(a) reports the length distribution of the SVs called by each tool on the HG007 sample. On HG007, the number of SVs reported by each tool ranges from 34,827 to 38,659 with `SVIM` reporting the lowest number of SVs and `SVDSS` reporting the highest number. Overall, all the tools report more insertions than deletions with shorter SVs (of length 100 *≤ bp*) being more frequent than longer ones. Moreover, all the tools show a clear peak at around 300bp suggesting a good capability in calling potential *Alu* mobile elements.

We also repeated the above experiment on HG007 using different aligners to test how SV callers are influenced by how reads are aligned. We tested all 6 callers in combination with `minimap2` [74] and `ngmlr` [75] (Supplementary Table S1). We also noticed that `SVDSS` significantly improves our ability to predict SVs in comparison to state-of-the-art approaches using `minimap2` mapper, while being one of top performer tools using `ngmlr` mapper (Supplementary Table S1).

Finally, we also investigated how read coverage affects SV calling performance. To this aim, we sub-sampled the HG007 sample (coverage 15x) down to 5x and 10x and we ran the 5 considered approaches on these two newly-created samples. Our `SVDSS` approach was also able to outperform other approaches using 10x sequencing coverage in all the metrics of interest (precision, recall, and F1, see (Figure 2(b) and Supplementary Table S2). When sample coverage is low (5x), `pbsv` achieves the highest recall (63.2%) at the expense of lower precision (58.6%) whereas other tools achieve similar high precision (ranging from 87.4% of `SVIM` to 92.9% of `SVDSS`) but low recall (ranging from 46.2% achieved by `SVDSS` to 51.6% achieved by `cuteSV`). As already pointed out in [72], `debreak` works poorly with low coverage samples. On the other hand, with higher coverages of 10x and 15x, `SVDSS` achieves the best precision and recall, outperforming other approaches.

### SVDSS improves SVs calling recall in hard-to-analyze regions

For further analysis, we divided the genome into two regions based on difficulty of calling variants based on GIAB’s definition of tiers [65]. Tier 1 accounts for nearly 86% of the genome spanning 2.51 Gbp, includes 50% or less of the total expected number of SVs, and is likely biased towards easy-to-call SVs ^3^. On the other hand, Tier 2 accounts for nearly 0.8% of the genome and consists of *∼* 6000 difficult-to-genotype sites. The remaining 13% of the genome mostly consists of centromeres, telomeres, and microsatellite regions (e.g., STRs) which are generally more difficult to genotype because of their repeat structure and due to the ambiguities of the reference genome. Because the high-quality assemblies that are the basis of our analysis include effectively complete genomes for each individual, we decided to extend Tier 2 to also include these regions (Extended Tier 2). This way, we are able to more thoroughly evaluate each methods accuracy across the entire human genome and we do not limit our analysis to easier-to-call regions (i.e., Tier 1). Supplementary Figure S3 provides a break-down of the tiers we considered in our analysis.

In this analysis, we considered the callsets produced by `SVDSS`, `cuteSV`, `pbsv`, `sniffles`, `debreak` and `SVIM` starting from `pbmm2` alignments. Table 1 reports the results of this analysis. Results on both tiers follow the same trend seen on full genome, with `SVDSS` managing to call more correct SVs without introducing many false calls. As expected, all tools achieve higher accuracy on Tier 1 regions, that are easier to analyze. Furthermore, we observed that the improvement between performance of `SVDSS` and other tools widens in the Extended Tier 2 regions of the genome (Table 1). Remarkably, on difficult-to-analyze regions (i.e., extended Tier 2), `SVDSS` achieves the highest recall, outperforming other callers by 15%, 14% and 4% on the HG002, HG007 and CHM13 samples, respectively.

To further provide evidence of correctness for true positive calls in these hard regions, we analyzed how these calls are shared among the tested callers using an upset plot [76]. Upset plots are an alternative to Venn diagrams that represents more conveniently the intersections of multiple sets. Figure 2(c) shows that out of the 10,333 total SVs in the truth set for HG007 (i.e., the `dipcall` callset), 3,720 (36%) of these SVs are correctly called by all the tested approaches whereas 2,399 (23%) are not detected by any tool. Remarkably, 739 SVs (7%) are detected only by our pipeline, partially explaining the higher recall it is able to achieve. `SVIM` has the second-highest number of specific calls at 130. Supplementary Figure S11 shows the distribution of `SVDSS`-specific vs `SVIM`-specific calls on chr1, chr2 and chr3 of the HG007 sample. SVDSS also detects the highest number of SVs that would have been exclusive to other tools, i.e., 172 (1.6%) calls are shared by `SVDSS` and `sniffles`, and 169 (1.6%) are shared between `SVDSS` and `pbsv`.

We manually investigated some of the SVs that are exclusively called by `SVDSS` and we discovered that most of such calls are SVs that exhibit two different alleles on the two haplotypes. These SVs account for heterozygous SVs with two non-reference alleles (as defined in [77]), i.e., SVs genotyped 1/2 (see two examples in Supplementary Figures S7 and S8) as well as pairs of close SVs whose alleles come from different haplotypes (see an example in Supplementary Figure S9). Other callers cannot always discern between the two haplotypes and heuristically call only one of the two alleles, inferring the length of alleles by combining the information coming from the two haplotypes. To further validate our claim, we considered each SV called only by our pipeline and we computed its distance to the closest SV. Out of a total of 792 SVs exclusively called by `SVDSS` and matching `dipcall` predictions using assembled genomes, 345 (44%) are located at exactly the same position as another called SV, hence are heterozygous SVs with two non-reference alleles, while 227 (29%) SVs are close (≤ 100bp) to another SV, and 107 (14%) SVs are too distant to another SV to be considered heterozygous events (Supplementary Figure S4). The `SVDSS` pipeline is able to more correctly manage these situations since it better discerns haplotypes supported by the input reads. Indeed, after clustering SV signatures by position, `SVDSS` splits each clusters in two subclusters, one per haplotype, hence allowing to call two different alleles - if needed.

### Hard-to-analyze regions harbor structural variants with clinical importance

To perform a more thorough analysis of the HG002 individual, we considered the CMRG (Challenging Medically Relevant Genes) callset provided in [52] and we evaluated callers’ accuracy against it. The CMRG callset consists of 250 SVs falling in 126 challenging and medically relevant genes that were excluded from the previously published GIAB benchmark [65] due to their complexity: compound heterozygous insertions, complex variants in segmental duplications, and long tandem repeats. The CMRG callset was created by diploid assembly of the haplotypes using hifiasm and then `dipcall`, proving one more time the effectiveness of assembly-based methods for detecting hard-to-analyze SVs, when well-curated assemblies are available.

As done previously, we computed the accuracy of `SVDSS` and the other 5 SV callers using `Truvari`. Out of the 250 SVs contained in the CMRG callset, `SVDSS` correctly called 232 SVs followed by `pbsv` (228) and `cuteSV` (225), `SVIM` (221), `debreak` (220), and `sniffles` (218). As shown in Figure 2(d), 5 SVs are exclusive to `SVDSS`, while 2 are missed exclusively by `SVDSS`: one was reported but with a length just under the evaluation threshold of `Truvari`, the other was missed due to being only detectable in clipped reads, which `SVDSS` does not consider by default. We then manually investigated the SVs that were exclusively called by `SVDSS`, discovering that all them exhibited two alleles, one per haplotype (i.e., heterozygous SVs with two non-reference alleles). This result confirms previous findings [52] that heterozygous insertions in tandem repeats are among the most challenging classes of SVs to discover with current methods.

Figure 3 shows one of the `SVDSS`-exclusive SVs, a double insertion inside the SLC27A5 gene on chromosome 19. Although the two haplotypes can be easily distinguished by visual inspection of adjacent heterozygous SNPs, the tested callers disagree on which allele to call. For instance only `SVDSS` calls two alleles of length 168bp and 224bp agreeing with the CMRG callset, whereas `pbsv` and `sniffles` report only one of the two (168bp). Surprisingly, `cuteSV`, `SVIM`, and `debreak` report a single allele of length 185bp, which does not match any of the evidence from read alignment. Additionally, we considered the portion of the high-quality HG002 assembly covering that locus (`chr19:58487900-58488500`) and we checked its alignment against the reference genome (Figure 3 and Supplementary Figure S5). Although the considered locus is in a repetitive region (as also proven by the noisiness of the dotplots shown in Supplementary Figure S5), the haplotype alignment confirm the presence of two alleles of different size.

### SVDSS has extremely low baseline error rate

Finally, we further investigate the lower bound on baseline false discovery rate of the proposed method by comparing the HiFi reads from CHM13 against the high-quality genome assembly built on the same samples (T2T [55]). Given the almost perfect T2T

CHM13 assembly produced using multiple orthogonal technologies, it is expected that an ideal SV caller would predict no SVs when comparing CHM13 reads against this assembly. Thus, we propose the following experiment to establish a lower bound on the baseline false discovery rate of different methods.

Ideally, our pipeline should generate zero SVs calls as no SFS should be extracted when querying CHM13 reads against the T2T assembly of the same sample. However, this will not be the case in practice due to the abundance of sequencing errors in particular from repetitive regions. Still, we expect the method to produce very few variant calls in this ideal scenario. The number of variants reported in such a scenario would also establish an empirical baseline for the method’s error-rate.

As a side-objective, we will also investigate the resulting SV calls to find if our method has discovered any true SVs missing from the T2T assembly. Due to the effectively homozygous nature of the CHM13 genome, any true variant discovered must be homozygous. However it is possible artifacts accumulated in the cell-line and actual heterozygosities in the genome may result in heterozygous SVs being reported.

We built the FMD index for v1.1 of the CHM13 assembly and extracted SFS from smoothed CHM13 PacBio HiFi data against this index. We then passed the SFS through the `SVDSS` pipeline for SV discovery. Our pipeline discovers a total of 102 SVs. For comparison, we repeated the above experiment with the other tools `pbsv`, `cuteSV`, `SVIM`, `debreak` and `sniffles`. Table 2 includes a summary of the result. We calculate the baseline False Discovery Rate for each tool as the number of calls it makes against T2T divided by the number of calls it makes against GRCh38. `SVDSS` has the lowest number of calls against the T2T assembly and also has the lowest baseline error rate.

**Table 2:** Comparison of baseline FDR rate of SVDSS with other methods. Number of SV calls against both the reference genome and the CHM13 assembly is included. Baseline FDR is calculated as division of first two columns for each tool. Number of known CHM13 heterozygous variants covered by each tool is included. The last two columns report the number of known CHM13 heterozygous (het) sites covered by each method and the precision of the method based on the number of covered heterzoygous sites.

| Tool | GRCh38 calls | T2T calls | Baseline FDR | Het Intersections | Het Precision |
| --- | --- | --- | --- | --- | --- |
| **SVDSS** | **23777** | **102** | **0.4%** | 13 | **12.7%** |
| cuteSV | 22654 | 667 | 2.94% | 23 | 3.4% |
| pbsv | 23707 | 616 | 2.59% | **28** | 4.5% |
| sniffles | 22680 | 314 | 1.38% | 22 | 7.0% |
| SVIM | 22176 | 948 | 4.27% | 29 | 3.0% |
| debreak | 23432 | 834 | 3.55% | 24 | 2.8% |

We further investigate if any of our calls are indeed true variants. The T2T project provides a list of known heterozygous sites on CHM13 ^4^ and 13 of our SV calls intersect these regions, suggesting that may be actual heterozygous alleles missing from the homozygous assembly. We also report the number of intersecting calls in Table 2 for every tool. `SVDSS` has the highest ratio of calls intersecting known heterozygous regions. We performed additional filtering of the calls using Merfin [78], a variant call polishing tool that filters VCF files based on whether the variants introduce *k*-mers not found in the sequencing reads. Only one of our calls passes Merfin’s filtering and we verify that the call seems to be a heterozygous site (Supplementary Figure S6).

In summary, `SVDSS` produces only 102 calls using CHM13 HiFi reads against the T2T CHM13 assembly, some of which may be actual true heterozygous variants. Furthermore, with our earlier experiments showing an average of 33,000 SV calls per sample, this amounts to a baseline error rate of less than 0.4% showing that `SVDSS` is robust to false detection of variants.

## 4 Discussion

We have introduced `SVDSS`, a hybrid method for SV discovery that combines advantages of different SV discovery approaches to achieve significant improvements in SV calling. A highlight of `SVDSS` is its much higher recall compared to other approaches in repeated regions of the genome (i.e., extended tier 2), and also its overall higher accuracy in particular in repetitive and traditionally hard-to-genotype regions of the genome. We also observed that reducing sequencing coverage impacts `SVDSS` less than other approaches. Thus `SVDSS` would provide much needed toolset to accurately predict SVs in low-coverage sequenced samples. Furthermore, utilizing the recent CHM13 assembly produced by T2T consortium, we could estimate baseline error rare for each methods and observed that `SVDSS` further has the lowest baseline error rate followed by `sniffles`.

While the availability of low-error long-read data enables more extensive variant discovery on new samples, SV discovery in repetitive regions of the genome such as STRs and microsatellites remains challenging but also hard to evaluate. This is evidenced by comparisons presented in this manuscript. Despite `SVDSS`’s significant performance improvements in repetitive regions, precision and recall in these regions are still lower than on the rest of the genome.

`SVDSS` currently supports the discovery of unbalanced structural variants, i.e. deletions and insertions, however as the underlying SFS signatures capture nearly all variation in the genome, a next step could be to extend the method to finding other classes of SVs such as inversions and duplications. Our current best technique for creating a SV truth sets (`dipcall`) does not evaluate inversions and duplications, yet a recent study [28] provides one of the first gold standards.

Throughout this work we highlight the importance of accurate benchmarks of SV calling methods. We evaluated `SVDSS` on a recent benchmark extensively curated over the HG002 sample [52] with the specific purpose of producing SVs occurring in genes of medical relevance, which are considered challenging for mapping-based and assembly-based SV prediction even from highly accurate long reads. This benchmark revealed that other methods fail to call heterozygous indels in highly homozygous regions or erroneous indels interpreted by a consensus approach. `SVDSS` is the only method able to discover 5 of such SVs in medically relevant gene regions.

## Footnotes

1 https://github.com/maxbachmann/rapidfuzz-cpp

2 https://github.com/ebiggers/libdeflate

3 https://ftp-trace.ncbi.nlm.nih.gov/giab/ftp/data/AshkenazimTrio/analysis/NIST_SVs_Integration_v0.6/README_SV_v0.6.txt

4 https://github.com/marbl/CHM13-issues

