## Supplementary Material for "Improved structural variant discovery in hard-to-call regions using sample-specific string detection from accurate long reads"

### A Supplementary Figures and Tables

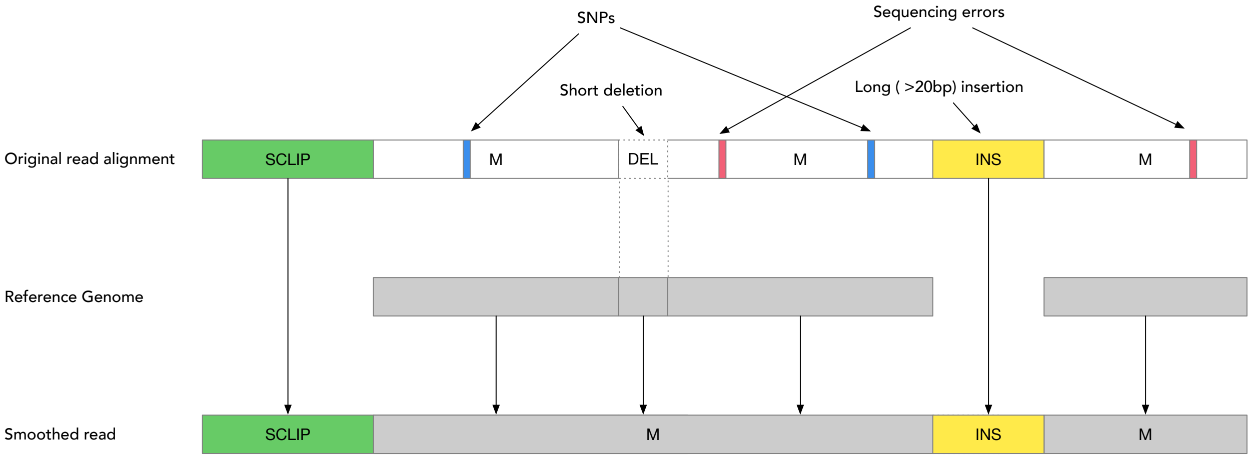


Figure S1: Illustration of the read smoothing algorithm. `M` alignment segments are smoothed from the reference genome, correcting SNPs (blue) and sequencing errors (red) in the process. Long indels are preserved while small ones ($< 20$bp) are removed. The small deletion is smoothed using the reference genome sequence while the large insertion (yellow) - potentially a SV - is carried over to the read. The soft-clipped section (green) is directly copied to the read.

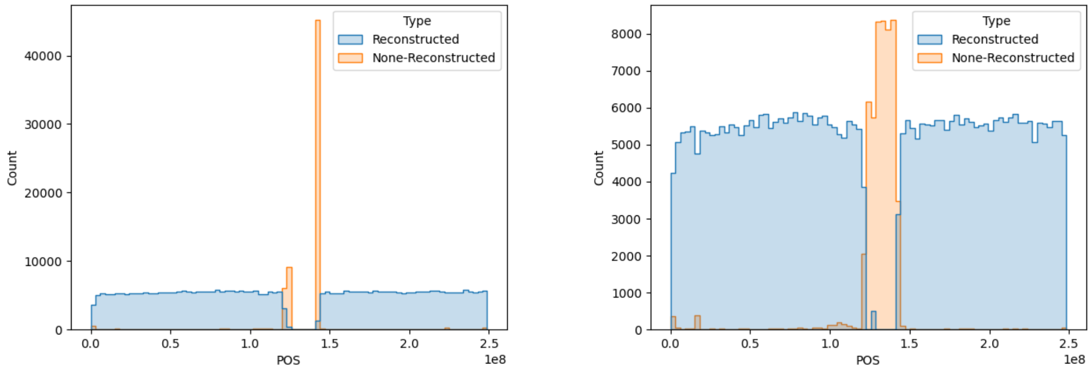


(a) GRCh38

(b) CHM13

Figure S2: Distribution of read mapping locations on chr1. Almost all non-smoothed reads originate from centromeres of CHM13, however almost none can be properly mapped the GRCh38, resulting in an alignment gap around the centromere.

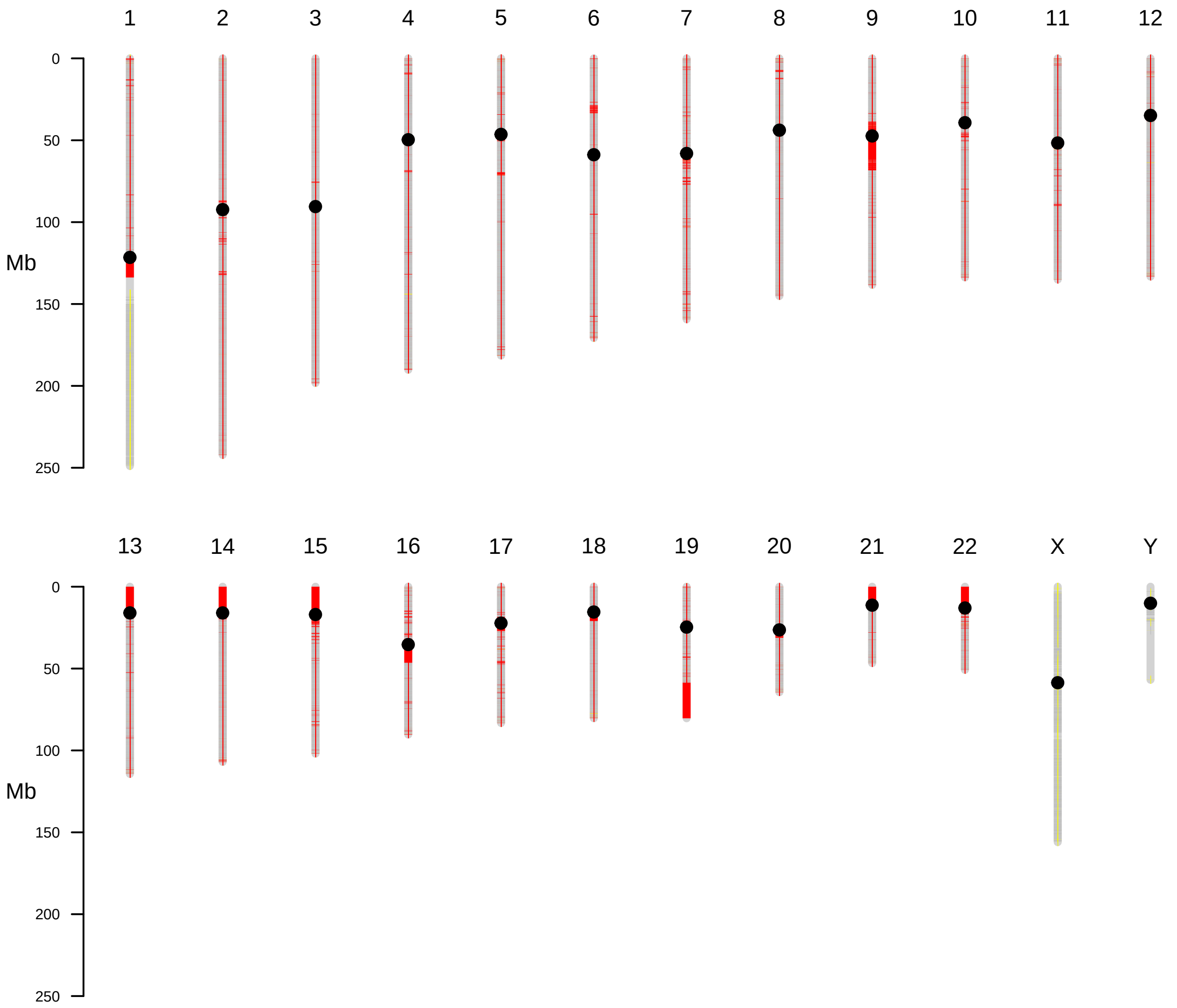


Figure S3: Tier 1 (grey areas) and extended tier 2 (red areas) regions in the reference genome GRCh38.

| Tool | pbmm2 P | pbmm2 R | pbmm2 F1 | minimap2 P | minimap2 R | minimap2 F1 | ngmlr P | ngmlr R | ngmlr F1 |
|---|---|---|---|---|---|---|---|---|---|
| SVDSS | **90.1** | **76.5** | **82.7** | **91.9** | **79.3** | **85.1** | **93.0** | 62.8 | 75.0 |
| cuteSV | 88.3 | 68.1 | 76.9 | 89.8 | 68.8 | 77.9 | 90.5 | 64.3 | 75.2 |
| pbsv | 84.9 | 68.6 | 75.9 | 85.0 | 68.7 | 76.0 | 84.9 | **67.9** | **75.5** |
| sniffles | 86.7 | 64.1 | 73.7 | 90.8 | 66.1 | 76.5 | 87.7 | 61.3 | 72.2 |
| SVIM | 84.9 | 64.7 | 73.4 | 87.2 | 65.7 | 74.9 | 81.9 | 64.4 | 72.1 |
| debreak | **90.1** | 64.2 | 75.0 | 91.3 | 64.8 | 75.8 | 86.0 | 60.8 | 71.2 |

Table S1: Comparison of performance of SVDSS and other methods when calling SVs on HG007 reads mapped with different aligners. Accuracy of each tool is reported in terms of Precision (P), Recall (R), and F-measure (F1). Results are whole-genome.

| Tool | 5x P | 5x R | 5x F1 | 10x P | 10x R | 10x F1 | 15x P | 15x R | 15x F1 |
|---|---|---|---|---|---|---|---|---|---|
| SVDSS | **92.9** | 46.2 | 61.7 | **91.6** | **70.3** | **79.5** | **90.1** | **76.5** | **82.7** |
| cuteSV | 92.0 | 51.6 | **66.1** | 90.5 | 65.2 | 75.8 | 88.3 | 68.1 | 76.9 |
| pbsv | 58.6 | **63.2** | 60.8 | 66.0 | 68.1 | 67.0 | 84.9 | 68.6 | 75.9 |
| sniffles | 91.4 | 46.9 | 62.0 | 89.7 | 60.0 | 71.9 | 86.7 | 64.1 | 73.7 |
| SVIM | 87.4 | 49.9 | 63.5 | 86.3 | 62.3 | 72.4 | 84.9 | 64.7 | 73.4 |
| debreak | 92.2 | 26.0 | 40.6 | 91.2 | 52.3 | 66.5 | **90.1** | 64.2 | 75.0 |

Table S2: Comparison of performance of SVDSS and other methods on HG007 at different coverages. Accuracy of each tool is reported in terms of Precision (P), Recall (R), and F-measure (F1). Results are genome-wide.

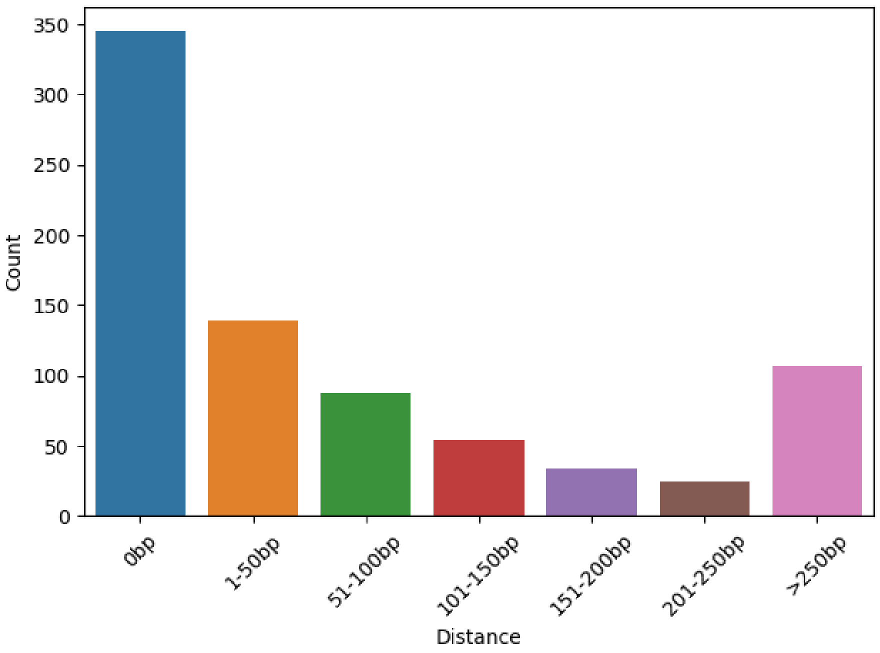


Figure S4: Bar plots showing the distance (in basepairs) of the SVs exclusively called by `SVDSS` and the closest SV (extended tier 2). A distance of 0bp means that the SV is a heterozygous non-reference SV (i.e., a SV with two alleles and genotyped 1/2).

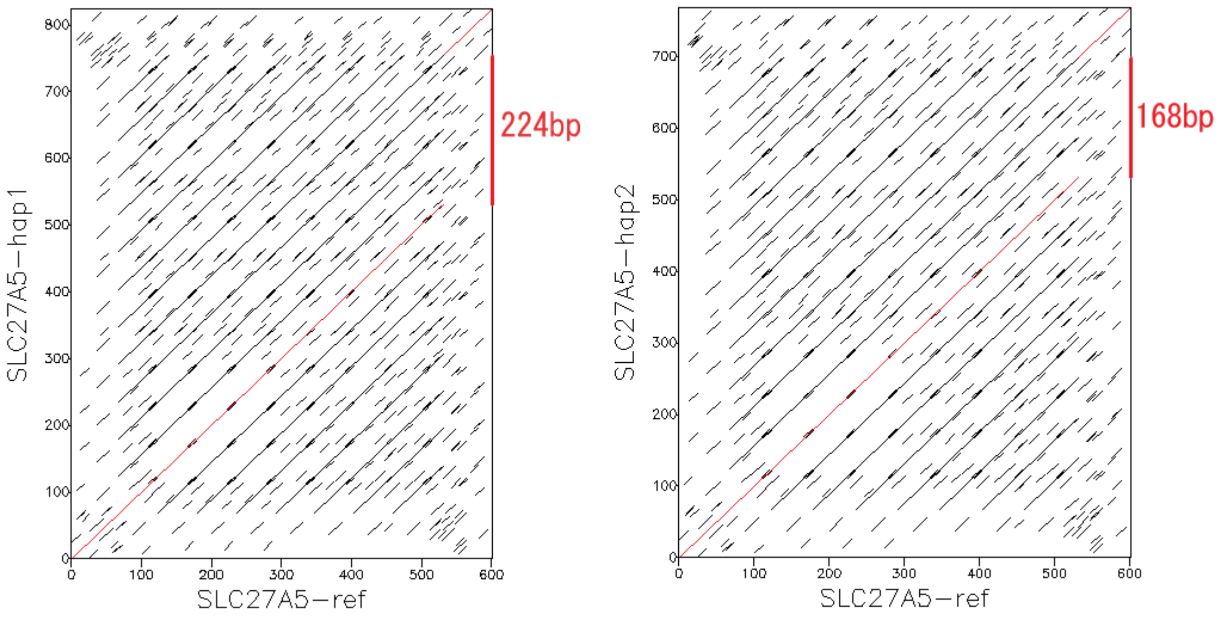


Figure S5: Dotplots of the alignment between the two HG002 high-quality haplotypes and the GRCh38 reference genome around the heterozygous SV falling in the medically-relevant gene SLC27A5 (locus: `chr19:58487900-58488500`).

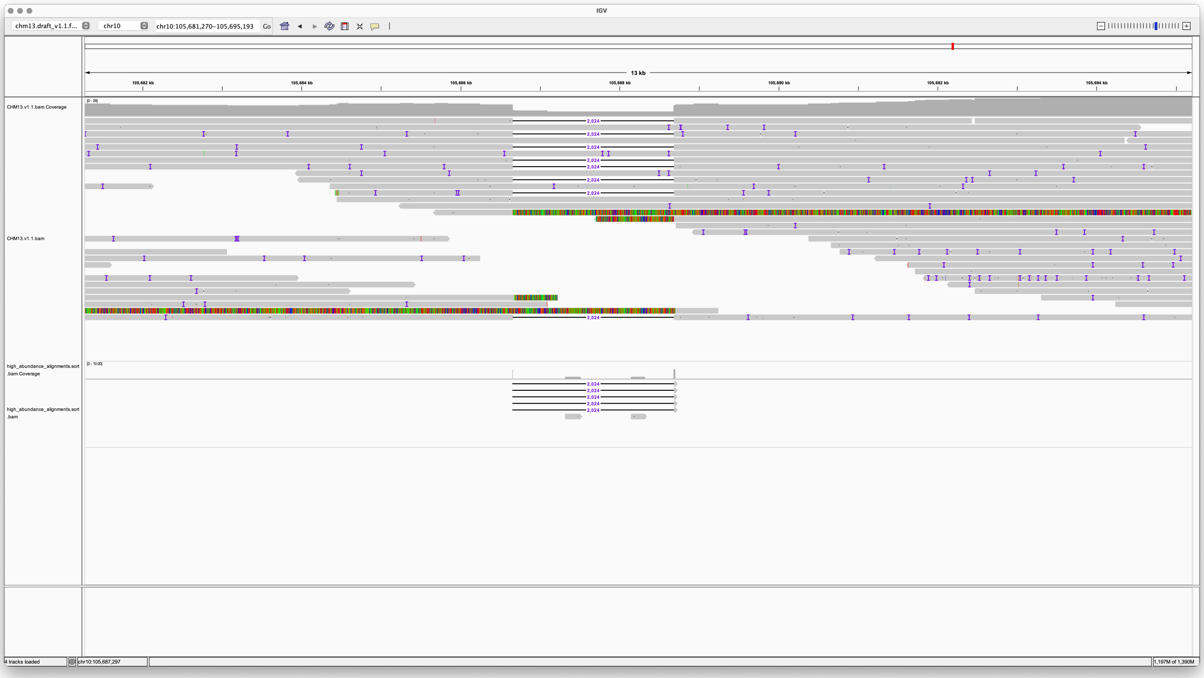


Figure S6: Example of heterozygous SV detected by our pipeline on CHM13 HiFi reads.

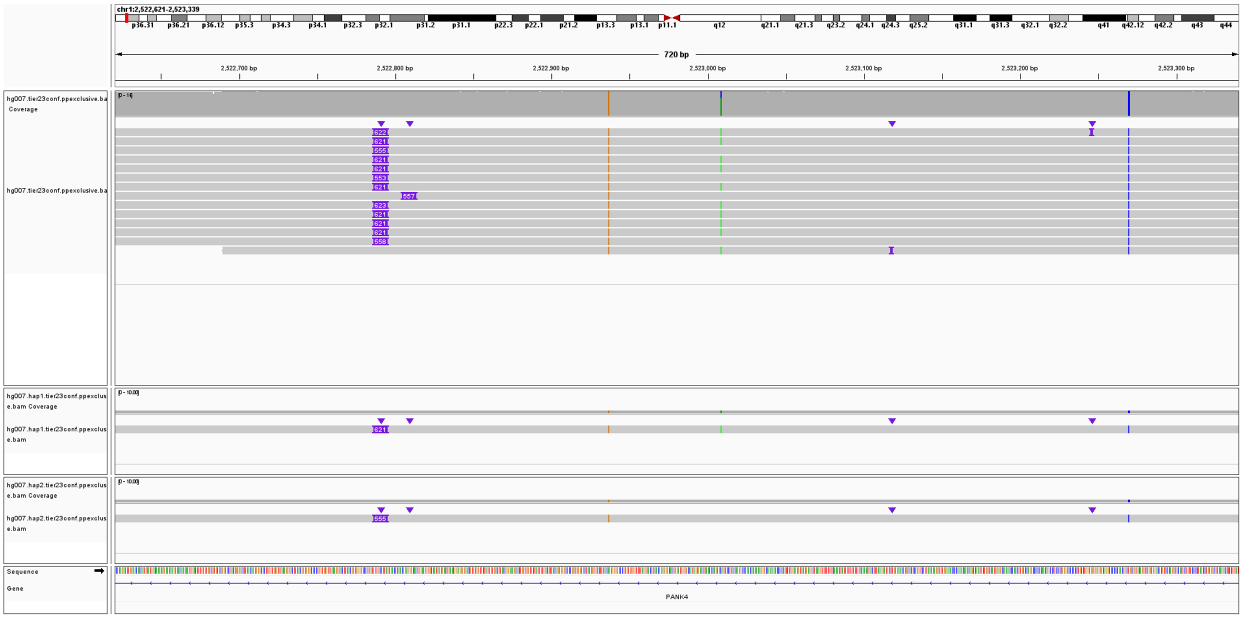

Figure S7: Heterozygous insertion in the HG007 sample (`chr1:2522791`). `dipcall` called two alleles of length 555 and 621. `SVDSS` agreed with `dipcall` correctly calling both alleles. `cuteSV`, `pbsv`, `sniffles`, `SVIM`, and `debreak`, instead, called just one allele of length 601, 621, 621, 602, and 601, respectively.

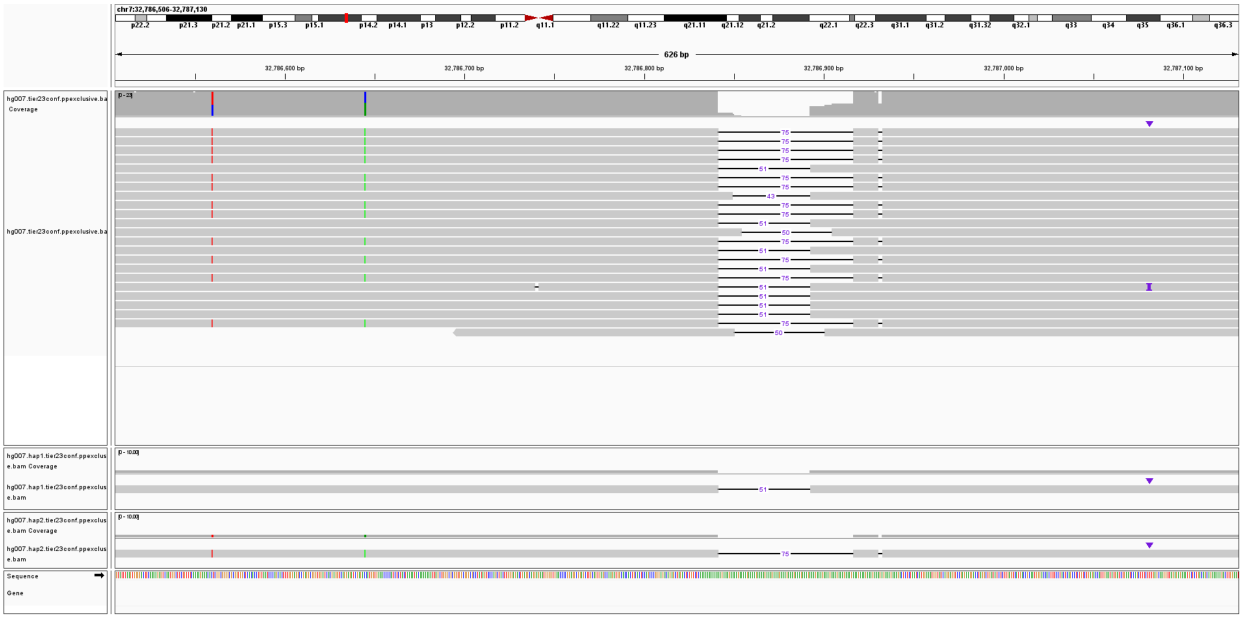

Figure S8: Heterozygous deletion in the HG007 sample (`chr7:32786841`). `dipcall` called two alleles of length 51 and 75. `SVDSS` agreed with `dipcall` correctly calling both alleles. `cuteSV`, `pbsv`, `sniffles`, `SVIM`, and `debreak`, instead, called just one allele of length 63, 75, 75, 63, and 63, respectively.

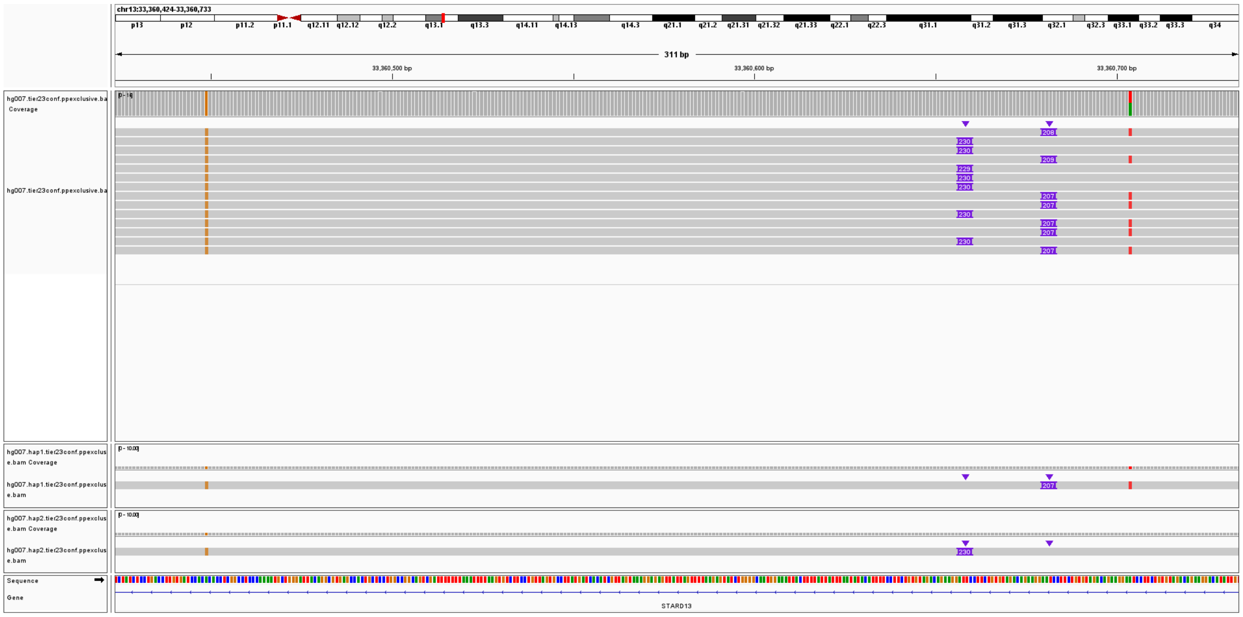


Figure S9: Two close insertion alleles in the HG007 sample (`chr13:33360658` and `chr13:33360681`). `dipcall` called two alleles of length 207 and 230. `SVDSS` agreed with `dipcall` correctly calling both alleles. `cuteSV`, `pbsv`, `sniffles`, `SVIM`, and `debreak`, instead, called just one allele of length 218, 230, 230, 218, and 218, respectively.

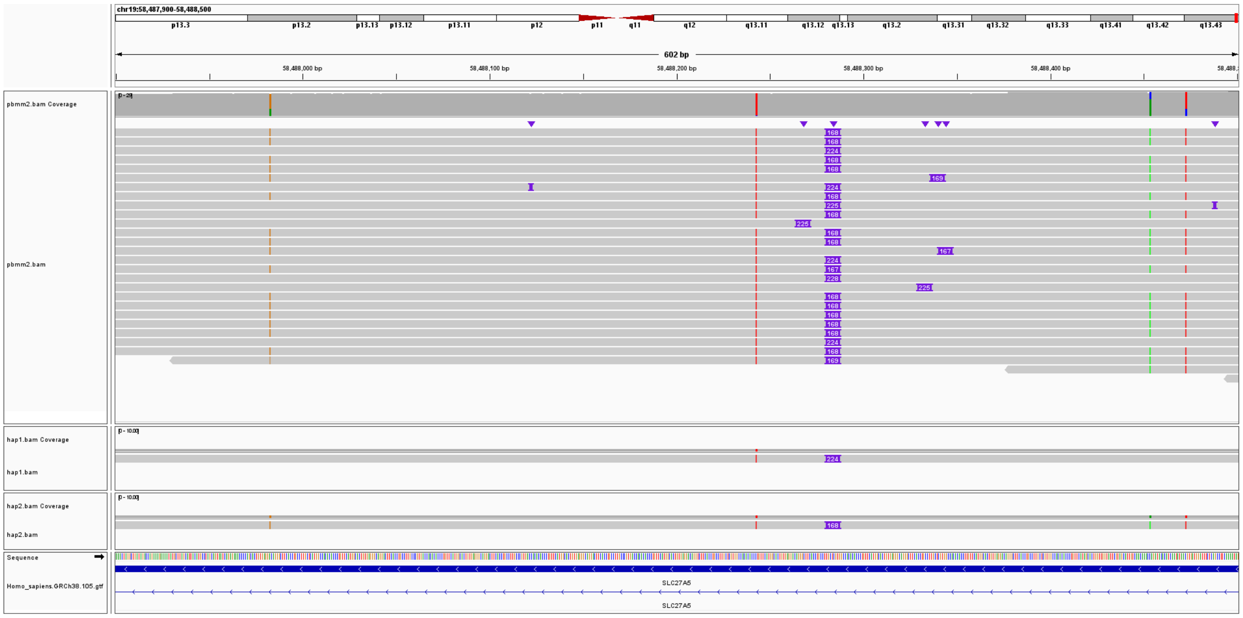


Figure S10: Full IGV image for the double insertion falling in the SLC27A5 medically-relevant gene.

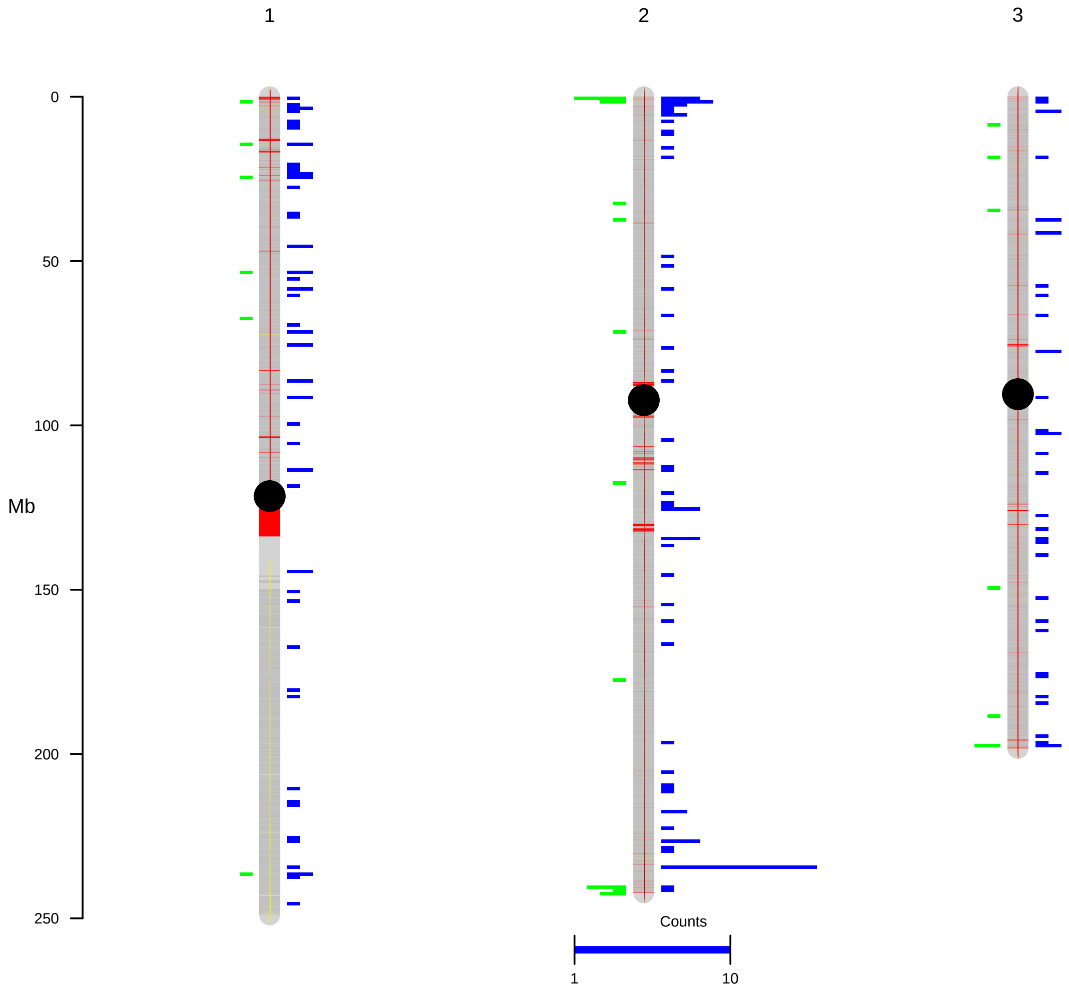


Figure S11: Comparison of `SVDSS`-specific and SVIM-specific Calls on Extended Tier 2 (red) on HG007. `SVDSS` (blue) has the highest number of specific calls on HG007 (739) while SVIM (green) has the second highest number of such calls (130).
